# Associations between Acoustic Features of Maternal Speech and Infants’ Emotion Regulation following a Social Stressor

**DOI:** 10.1101/2021.07.02.450379

**Authors:** Jacek Kolacz, Elizabeth B. daSilva, Gregory F. Lewis, Bennett I. Bertenthal, Stephen W. Porges

## Abstract

Caregiver voices may provide cues to mobilize or calm infants. This study examined whether maternal prosody predicted changes in infants’ biobehavioral state during the Still Face, a stressor in which the mother withdraws and reinstates social engagement. Ninety-four dyads participated in the study (infant age 4-8 months). Infants’ heart rate and respiratory sinus arrhythmia (measuring cardiac vagal tone) were derived from an electrocardiogram (ECG). Infants’ behavioral distress was measured by negative vocalizations, facial expressions, and gaze aversion. Mothers’ vocalizations were measured with spectral analysis and spectro-temporal modulation using a two-dimensional fast Fourier transformation of the audio spectrogram. High values on the maternal prosody composite were associated with decreases in infants’ heart rate (β=-.26, 95% CI: [-.46, -.05]) and behavioral distress (β=-.20, 95% CI: [-.38, -.02]), and increases in cardiac vagal tone in infants whose vagal tone was low during the stressor (1 SD below mean β=.39, 95% CI: [.06, .73]). High infant heart rate predicted increases in the maternal prosody composite (β=.18, 95% CI: [.03, .33]). These results suggest specific vocal acoustic features of speech that are relevant for regulating infants’ biobehavioral state and demonstrate mother-infant bi-directional dynamics.

---

Infants’ biobehavioral regulation relies heavily on interactions with caregivers, which set the stage for the development of increasing self-regulation with age (Feldman, 2007a; Tronick, 1989). The acoustic features of caregiver voices may be a key influence on infants’ states (Spinelli, Fasolo, & Mesman, 2017) and these features are emphasized in multiple theoretical models of infant-caregiver interactions (Feldman, 2007a; Tronick, 1989; Welch & Ludwig, 2017). Prior research shows that caregiver vocalization occurrence, contingency, use of infant-directed speech and singing, and emotional content as rated by judges all reflect salient aspects of caregiver-infant interactions (Goldstein, Schwade, & Bornstein, 2009; Shenfield, Trehub, & Nakata, 2003). However, specific prosodic acoustic features – those which form the foundation of the emotional and affective components of vocalizations – have received little attention. In this study, we use predictions based on the Polyvagal Theory (Porges, 1995, 2001, 2007), an evolutionary neurophysiological framework, to inform the selection of frequency and spectro-temporal acoustic features that reflect prosodic features in humans. We then examine how maternal prosodic features facilitate infants’ biobehavioral regulation following a social stressor – the Still Face Paradigm (SFP; Tronick, Als, Adamson, Wise, & Brazelton, 1978).

## Maternal Vocal Communication with Infants

Infants respond to their mothers’ voices in utero beginning at 32 weeks gestational age, as demonstrated by changes in heart rate (HR) (Kisilevsky et al., 2009). As a result of these in utero experiences, newborns may develop preferences to listen to their native language and to their mother’s voice (Decasper & Fifer, 1980; Moon, Cooper, & Fifer, 1993). Over the first few months of life, mothers and their infants establish patterns of mutual vocal responsiveness, (Beebe, Alson, Jaffe, Feldstein, & Crown, 1988; Lavelli & Fogel, 2013), including contingently responding to each other’s vocal cues (Bigelow & Rochat, 2006). Maternal speech toward the infant also increases in both amount and complexity during this time (Henning, Striano, & Lieven, 2005).

Infants are sensitive to the prosody of their mothers’ voices and the communicative intent it conveys (Fernald, 1989), preferring to listen to prosodic infant-directed speech’ compared to a more monotonic adult-like speech (Cooper & Aslin, 1990), especially the infant-directed speech of their own mother (Cooper, Abraham, Berman, & Staska, 1997). Infant-directed speech is characterized by exaggerated modulation of pitch and slower rhythm and tempo (Fernald & Kuhl, 1987) and conveys emotional information to infants (Trainor, Austin, & Desjardins, 2000). Using near infrared spectroscopy in the temporal cortex, researchers have found that between 4 to 7 months of age, infants’ neural responses to emotional voices become enhanced relative to neutral voices (Grossmann, Oberecker, Koch, & Friederici, 2010), suggesting sensitivity to emotional information in the voice in early infancy.

Mothers may especially rely on vocalizations to not only interact with their infants, but also to help regulate their infants’ affect and behavioral reactivity, particularly during times of stress. Research has shown that mothers’ singing modulates activity of the Hypothalamic-Pituitary-Adrenal axis (Shenfield et al., 2003), promotes infant behavioral and physiological regulation of the sympathetic nervous system (Cirelli, Jurewicz, & Trehub, 2020; Cirelli & Trehub, 2020) and reduces incidence of infantile colic in the first two months after birth (Persico et al., 2017). However, few studies have examined the the acoustic features of speech as a source of biobehavioral regulation for infants.

## The Polyvagal Theory and Acoustic Regulation of State

Polyvagal theory describes the developmental and evolutionary transitions in the neural pathways that regulate biobehavioral state, providing a foundation for infant-caregiver emotional co-regulation (Porges, 1995, 2001, 2007). In mammals, myelinated vagal pathways provide parasympathetic influence on the heart. These neural pathways efficiently reduce heart rate to promote calming and are bidirectionally linked with the brainstem areas controlling the structures involved in social engagement (e.g., facial expression, gaze, listening, and vocalizations; Porges, 2001). This integrated social engagement system continues to develop through the early years of life, supporting increasing capacities for interpersonal co-regulation that promotes calm states for affiliative interactions (Porges & Furman, 2011).

The Polyvagal Theory also proposes that mammalian biobehavioral state regulation is influenced by the frequency band and spectro-temporal modulation of auditory cues (Kolacz et al., 2018; Porges & Lewis, 2010). The mammalian frequency band of perceptual advantage (optimal hearing range) is influenced by the physics of the middle ear, with the human adult band ranging from about 500-4000Hz (Kolacz et al., 2018; Porges & Lewis, 2010). The theory proposes that this band co-evolved with the neural regulation of the physical structures that influence the acoustic features of vocalizations (e.g., larynx and pharynx) and that these structures are functionally integrated with brainstem regions that connect the autonomic nervous system with higher brain regions (Porges, 1995; 2001; 2007). This integration provides an acoustic channel for physiological co-regulation, informing the hypotheses that biobehavioral safety cues may be carried in the mammalian frequency band of perceptual advantage and in the frequency modulation characteristics. Frequency and amplitude modulation are common features of social affiliative communication across a range of mammalian species while defensive or distress calls tend to be more monotone (Brudzynski, 2007; see review in Porges & Lewis, 2010). Taken together, safety cues defined by frequency band and modulation characteristics may promote parasympathetic regulation and reduce biobehavioral arousal to support calm engagement for affiliative social interactions. Conversely, danger cues may be communicated outside the frequency band of perceptual advantage, such as within high frequencies that can be used for distress calls, which may also have more monotone features with limited spectro-temporal modulation (Porges & Lewis, 2010). These danger and life threatening cues would promote distress, mobilization, or behavioral shut down. Thus, maternal prosodic features that promote calming in infants may be marked by relatively strong power in the frequency band of perceptual advantage, relatively little power in high frequencies, and high levels of spectro-temporal modulation.

## The Still Face as a Social Stressor

Acoustic features of the mother’s voices are thought to be especially important for soothing infants’ distress or co-regulating after a disruption in contingency (Feldman, 2007b; Gianino & Tronick, 1988; Tronick, 1989). The Still-Face Paradigm (Tronick et al., 1978) is an experimental simulation of the interruption of the infant-parent interaction. After a period of face-to-face dyadic interaction, the parent is instructed to terminate the interaction and gaze at the infant with an inexpressive, unresponsive face known as the ‘still-face’ (SF). After this disruption which typically lasts 1-2 minutes, the social interaction resumes in the ‘reunion’ phase. Infants tend to significantly increase negative affect from face-to-face play to the still-face, and then decrease negative affect once maternal interaction resumes in reunion (e.g. Mesman, van IJzendoorn, & Bakermans-Kranenburg, 2009). There is often, however, a carry-over effect such that negative affect in reunion doesn’t completely rebound to comparable levels observed in face-to-face play prior to the still-face perturbation (Weinberg & Tronick, 1996).

These behavioral changes are accompanied by changes in physiological state across phases: infants typically increase heart rate and decrease cardiac vagal tone in response to the still-face (suggesting a decrease in safety-related parasympathetic regulation), followed by subsequent decreases in heart rate and increases in vagal tone during reunion (Jones-Mason, Alkon, Coccia, & Bush, 2018).

To our knowledge, one previous study has examined the role of maternal speech prosody in regulating infants’ state in the context of a stressor (Spinelli & Mesman, 2018). When comparing measures of maternal fundamental frequency variability (a marker of prosodic modulation of voice) and mean (a measure of pitch), only the former was a predictor of lower infant negative affect after the Still Face (though the effect was moderated by maternal sensitivity, see Spinelli & Mesman, 2018). These results support the particular importance of prosodic modulation for infant affective regulation, though the interpretation of these results in the context of dynamic regulation is limited due to the measurement of maternal prosody during baseline play but not following the stressor.

The current study extends previous work by focusing on maternal prosody during the reunion, to target a component of the regulatory processes that occur after the stressor. The reunion phase poses its own regulatory challenge for infant-mother dyads: for the mother, the challenge is to re-engage and soothe her infant following the still-face perturbation. For infants, the challenge is to reconcile the mother’s sudden return to interaction with her previous bout of non-interaction (Weinberg & Tronick, 1996). As such, the reunion phase is ripe for providing valuable information regarding the quality of dyadic co-regulation (Cohn, 2003).

The current study examines infants’ physiological regulation in relation to maternal prosody following the social stressor of the Still Face Paradigm. We hypothesized that high maternal prosody – indexed by strong mid frequency power, low leves of high frequency power, and spectro-temporal variability – would predict a) decreases in infant heart rate (HR), a measure of overall cardiac output, b) increases in infant cardiac vagal tone, reflecting greater parasympathetic activity, and c) decreases in behavioral distress following the Still Face. In addition, we conducted exploratory analyses of the moderation of infants’ biobehavioral state prior to re-engagement on the associations between maternal prosody and infant state after the stressor. These were conducted to examine whether high or low values of behavioral distress or physiological mobilization were differentially predictive of changes associated with maternal acoustic features. Finally, we conducted an exploratory cross-lagged panel analysis with maternal prosody and infants’ biobehavioral variables to examine the directionality of effects within the dyads.

## Method

### Participants

Ninety-six mothers and their 4-to 8-month-old infants were recruited from the local community of a Midwestern college town as part of a larger longitudinal study of children’s self-regulation. Infants were eligible for the study if they were born full term (at least 37 weeks gestational age at birth). Parents’ contact information was obtained through public birth records, voluntary sign-up at community events, and referral by other families. Infants were given a small toy for participation. The [BLINDED FOR PEER REVIEW] Institutional Review Board (IRB) approved all protocols for the study. Overall, the sample reflects the local county demographics, which is 83.4% White, 7.0% Asian, 3.6% Black, 3.5% Hispanic and 2.5% Multi-racial or other according to the most recent census (US Census Bureau, 2018). Two infants were excluded due to camera malfunction during the session, resultingin a final sample of 94 infants (age M= 5.18; SD = 1.31; 49 males).

### Procedure

Lab visits were scheduled at a time when the infant was normally awake and alert. Upon arrival to the laboratory, a female experimenter explained the procedure and obtained informed consent from the mother. Mothers were seated approximately 30 cm away facing their infants, who were placed in an infant seat (Figure 1). Two cameras were positioned unobtrusively in the corners of the room to record audio and video; one facing the child and the other facing the mother.

**Figure 1.**
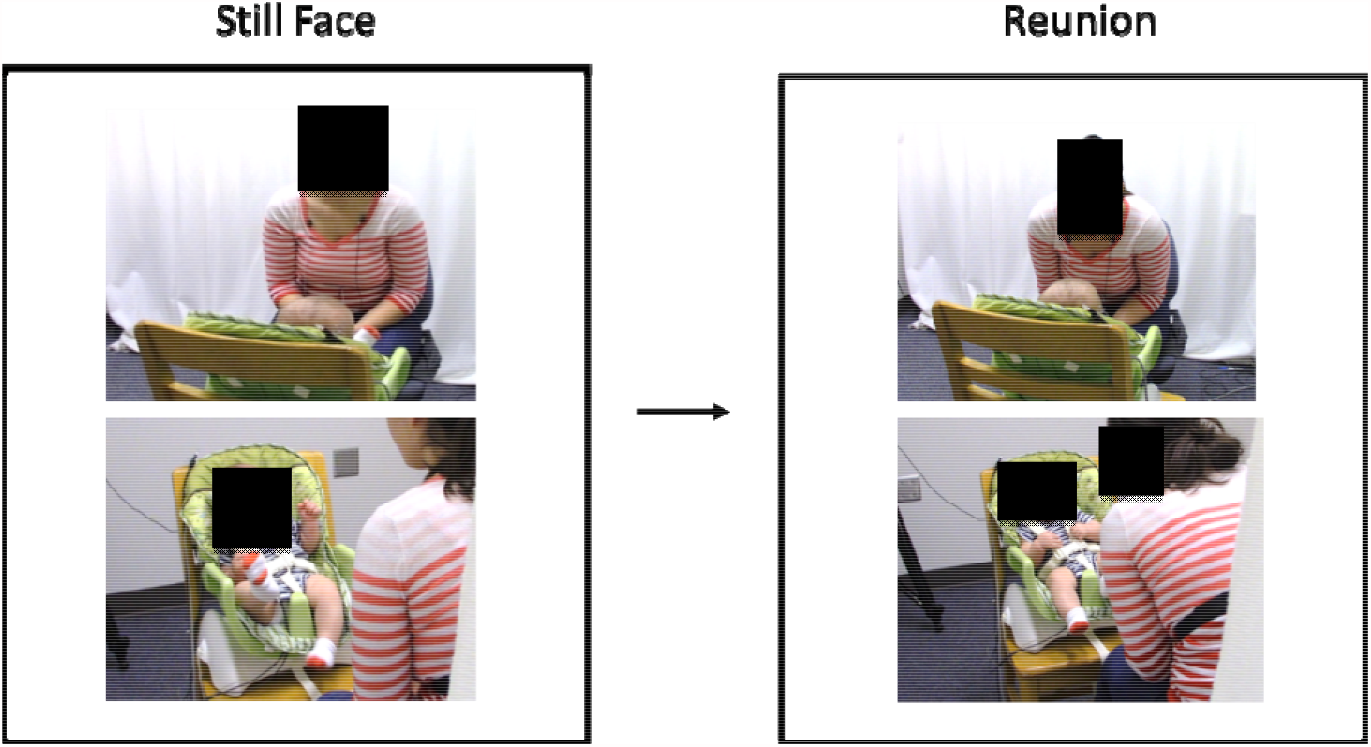
Caregivers and infants were seated across from each other during the interaction. Cameras on both sides of the room recorded both interaction partners’ behavior. Recordings from the microphone mounted within the camera facing the mother was used to assess acoustic features of the caregivers’ vocalizations. During a 2-minute “Still Face” episode (Tronick et al., 1978), caregivers were not allowed to make facial expressions, vocalize, or touch the infant. During the “Reunion” phase, caregivers could interact with infants as they normally would. Changes in infant physiological regulation and behavioral distress were tracked from the Still Face episode to the end of reunion. Caregivers’ vocal features were assessed at beginning of the reunion period as caregivers and infants re-established contingent interactions.

To monitor cardiac physiology, three disposable Ag/AgCl electrodes were placed on the infants’ chest in an inverted triangle pattern. These were connected to electrode leads that were hidden under the infant’s clothing and seat cover. Electrocardiogram signals were recorded at a 1000Hz sampling rate using the Biopac MP150 system (Biopac Systems Inc., Goleta, CA).

Mothers’ ECG was also collected but not used for this analysis. Following set-up and explanation of procedures to the mother, the experimenter monitored the dyad from behind a large curtain using a third camera.

At the start of the protocol, an infant ECG baseline measure was collected for 30 seconds while the mother completed a questionnaire. Mother and infant then took part in the Face-to-Face Still Face paradigm (Tronick et al., 1978) consisting of three sequential two-minute phases that were verbally cued by the experimenter. First, in the *face-to-face play* phase, mothers were instructed to interact normally with their infants. Mothers were asked not to touch their infants, as this might interfere with heart rate recordings (Feldman, Singer, & Zagoory, 2010). During the *still-face* phase, mothers were instructed to pose a ‘still-face’— looking at their infants’ forehead with a neutral expression while keeping their face and body still. After two minutes, mothers were cued to resume normal social interactions (*reunion* phase). If at any time infants became too distressed (i.e. hard crying for more than 10 seconds), the session was terminated early at the discretion of the mother or experimenter. Mothers were not provided any specific instructions about their vocalizations.

#### Physiological Data Quantification

Infants’ heart rate and respiratory sinus arrhythmia (RSA) were extracted from the ECG signal. The digitized ECG data were analyzed using a peak detection algorithm to capture the time interval between sequential R-waves to the nearest ms. CardioEdit software (Brain Body Center, University of Illinois at Chicago) was used for visual inspection of data and correction of artifacts caused by movement and missed beats. Respiratory sinus arrhythmia (RSA) was measured using the method developed by Porges & Bohrer (Lewis, Furman, McCool, & Porges, 2012; Porges & Bohrer, 1990) and implemented in the CardioBatch+ software (2015). First, heart period values were resampled into equal 200ms time intervals. Then, aperiodic baselines and slow oscillations were removed using a moving polynomial filter (3^rd^ order, 21 point, 4.2 sec duration) and the data were band-passed in the frequency range of spontaneous breathing for infants (0.3-1.3 Hz). The resulting variance was natural log transformed to reduce skewness, generating RSA estimates in ln(ms)^2^ units. RSA and heart rate were calculated in 30-sec epochs and averaged over the phases of interest.

#### Maternal prosody analysis

Maternal audio files were extracted from the recorded videos. Audio was recorded with the mounted on-camera condenser microphone with an upper frequency bound of 16 kHz. Mothers’ voiced speech was manually isolated – removing singing, whispering, infant noises, and extended stretches of silence - using Audacity software (Audacity Team, 2018). Speech was isolated because of its predominance as a mode of vocalization. All mothers used speech during reunion, and its use accounted for 85.9% of total maternal vocalizations. Though all vocalizations are likely to contribute to infant biobehavioral regulation, each type of vocalization also has unique acoustic properties. For instance, prior research shows that maternal song can maintain or reduce infant sympathetic activity, depending on whether singing is performed in a soothing or playful way (Cirelli et al., 2020). However, infant-directed singing is less variable in pitch, dynamics, and timing (e.g. Corbeil, Trehub, & Peretz, 2013) and these types of categorical acoustic differences may cause confounding. Given that non-speech vocalizations were rare (e.g., 16% of mothers sang and singing constituted less than 10% of total vocalizations), they were excluded from the main analysis in order to focus on the most predominant and relatively homogenous acoustic signal.

Prior to analysis, recordings were modified by a high-pass Chebyshev II filter (cut-off frequency = 180 Hz, order = 65) to attenuate low-frequency environmental noise. The resulting files were then analyzed for relative power in the high (>5000 Hz) and mid (500-5000 Hz) bands using a Fast Fourier transformation and spectro-temporal modulation as described below. Because middle ear bones are smaller at early maturational stages, infants’ frequency bands of advantage extend to higher frequencies than those of adults. Thus it would be expected that the threshold between their safety-related acoustic channel and high frequency danger-signaling channel would be higher than adults. For the purposes of this paper, we tentatively set that threshold at 5000Hz, though future research is needed to examine how high this threshold extends.

Spectro-temporal modulation was analyzed using a modified version of the modulation power spectrum Matlab package developed by Singh & Theunissen (2003). The modulation power spectrum (MPS) is a two-dimensional Fast Fourier transformation that decomposes the time-varying acoustic signals of a spectrogram (Figure 2) into a two-axis space of spectral (frequency) and temporal (time-related) modulation. These joint spectro-temporal representations have been found to characterize auditory neurons’ higher level processing of natural sounds (Theunissen & Elie, 2014) and the organization of the auditory cortex including feature detectors that correspond to specific spectro-temporal patterns (Norman-Haignere, Kanwisher, & McDermott, 2015). The MPS was applied to 1.5-second sequential time slices with 50% overlap and averaged over the length of the audio file. Time windows were selected to maximize the frequency resolution of the spectrogram without collapsing multiple utterances into a single window. Frequency-specific background noise was estimated for each recording by averaging the spectral density across the 5% lowest intensity time windows of these spectrograms. The difference between the spectrogram and this estimated level of background noise was calculated and used to derive the MPS metrics. This second de-noising step mitigated differences in the amplitude and frequency content of the remaining environmental noise, which could vary from recording to recording due to multiple factors including individual differences in maternal speaking volume. The correction was applied at the level of the spectrogram, not the waveform. Thus, the 2D-FFT was applied to segments of a spectrogram that reflected ‘energy above background noise’, eliminating a confounding source of unmodulated sound energy (e.g., the mechanical noise of the ventilation system). Modulation depth was operationalized as the square root of the ratio between power at origin (0, 0), reflecting unmodulated vocal sound, to power in the rest of the spectrum (Singh & Theunissen, 2003).

**Figure 2.**
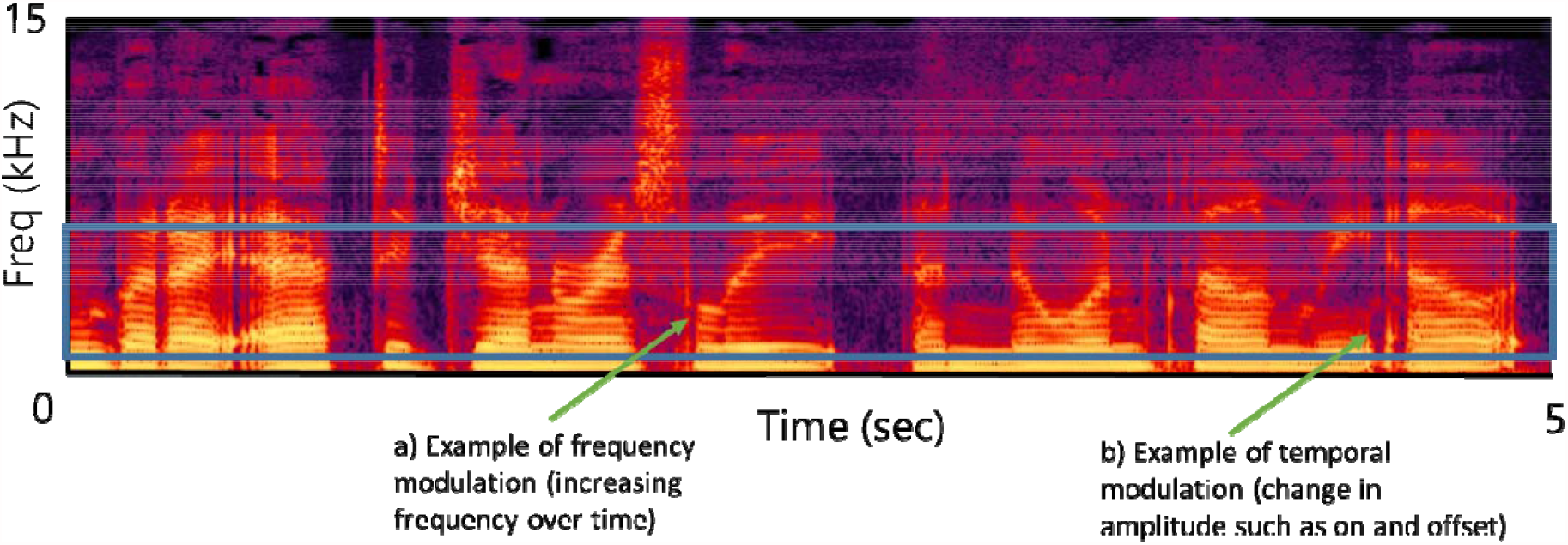
A spectrogram of female adult speech, with time on the x axis and frequency on the y. Lighter colors represent higher acoustic power. Frequencies within the blue box span 500-5000 Hz (mid frequency power). Frequency modulation occurs when a tone changes in frequency over time (a) and temporal modulation occurs with amplitude changes in amplitude (e.g., onset and offset; b). Relative power of maternal vocalization frequency bands was measured using an FFT of the waveform. Maternal spectro-temporal modulation was assessed by modulation power spectrum (MPS), a 2□dimensional fast fourier transformation (FFT) applied to the spectrogram (Singh & Theunissen, 2003).

Based on theoretical predictions and the properties of the middle ear (see introduction), maternal prosody composite values were created so that high prosody scores would reflect strong mid frequency power, low levels of high frequency power, and high spectro-temporal modulation. First, raw variables were transformed to z scores. The sign of high frequency power was reversed so that lower power would correspond with higher composite values (i.e., greater prosodic features). The mean of the resulting mid frequency power, high frequency power (reversed), and modulation depth z scores were used as a prosody composite value.

#### Data Coding

##### Observation codes

Infants’ facial expressions, vocalizations, and eye gaze were coded using Observer software (Noldus Information Technology Inc., Leesburg, VA). Measures of facial distress included tension between the eyebrows, grimace, tension in lips (lip pursing), and turning down of the corners of the mouth (Mattson et al., 2013). Distressed facial expressions are often accompanied by negative vocalizations ranging from subtle perturbations such as fussing and whimpering to more intense negative vocalizations, namely crying (Haley & Stansbury, 2003; Weinberg & Tronick, 1996). Negative vocalizations included fusses, whines, and cries.

Eye gaze was coded as directed towards the social partner’s face or body, at the social partner’s hands or at an object, or gaze aversion away from social partner, including looking anywhere around the room or at one’s own body. Behavior was continuously coded every second; codes within each category were mutually exclusive and exhaustive (for full list see Table S1).

##### Reliability

Reliability was calculated for 15% percent of all videos. In the Observer program, codes were evaluated for code matching and duration. Raters achieved the following percent agreement for infants’ behaviors: vocalizations (81%), facial expressions (86%), and gaze (76%). To control for chance agreement, kappa scores were also calculated. Raters achieved substantial agreement (Landis & Koch, 1977) across all categories: vocalizations (K= 0.64), facial expressions (K= 0.81) and gaze (K= 0.68).

##### Behavioral distress

A composite measure of distress was created based upon the amount of time infants exhibited negative facial expressions, negative vocalizations, and gaze aversion, as these behaviors tend to co-occur (Pratt, Singer, Kanat-Maymon, & Feldman, 2015; Stifter & Moyer, 1991). In order to credit infants for expressing more than one negative behavior at a time infants received a score of 0 to 3 for every second they were observed. If infants expressed only one behavior, they received a score of “1”; two behaviors resulted in a score of “2”, and three behaviors resulted in a score of “3”. Scores were summed across 30 second quartiles, which was then divided by the duration of the quartile (typically 30 sec) to compute a composite distress score for each quartile. Final scores were then divided by 3 so that they ranged between 0-1. In cases in which the full 30-second quartile was disrupted (e.g., by termination of the protocol by caregiver or researcher due to extreme infant distress), missing data were imputed using the previous quartile’s value. Behavioral distress measures were transformed using ordered quantile normalization to reduce bias caused by skewness in the raw metric (Peterson & Cavanaugh, 2019).

## Data Analysis

Models were fit and visualized using R 4.0.1 (R Core Team, 2020). For regression analysis, mothers’ prosody variables were calculated during the first minute of reunion to capture vocal features during the reorganization in transition from Still Face to congintent responding. On average, mothers’ isolated vocalizations composed 54% of total audio after the Still Face when other sounds and silences were excluded (for 1-minute segments of audio: M = 32.45 sec, SD = 14.74).

To examine infants’ changes in biobehavioral state, raw change in heart rate, RSA, and behavioral distress were calculated as outcome variables. These were calculated as the difference between final minute of reunion and still face values so that negative values represent decreases, positive values represent increases, and 0 represents no change. Linear regression models were used to predict infants’ biobehavioral changes from the Still Face to the second minute of reunion as a function of mothers’ vocal acoustic features during the first minute of reunion. Follow up analyses were conducted to examine interactions between prosodic features and infants’ state during the Still Face episode.

Cross-lagged panel models were used to examine temporal dynamics of the maternal prosody composite and infant state measures. These models were fit using the lavaan R package (Rosseel, 2012) with the maximum likelihood estimator, a method that permits use of all observations even when missing data are present. Model fit was evaluated using the root mean squared error of approximation (RMSEA; Steiger, 1990), the Tucker-Lewis Index (TLI; Tucker & Lewis, 1973); and the Comparative Fit Index (CFI; Bentler, 1990). Good model fit was assessed by CFI and TLI values near or above .95, as recommended by Hu & Bentler (1999).

## Results

### Univariate and Bivariate Descriptive Statistics

Infants’ age was not correlated with any prosody variables (all p > .20). As described above, only maternal speech – which comprised 85.86% of all maternal vocalizations - was used for main analysis (additional analyses using all vocalizations and singing are reported in Supplemental Material). Univariate descriptive statistics are presented in Table 1. On average, infants’ behavioral distress and heart rate decreased from Still Face to the second minute of reunion, while infants’ RSA increased (Table 2).

**Table 1.** Descriptive Statistics.

| **Variable** | **n** | **Mean** | **SD** | **Median** | **Min** | **Max** |
| --- | --- | --- | --- | --- | --- | --- |
| *Maternal Prosody* |  |  |  |  |  |  |
| Maternal Prosody Composite Reunion Min 1 | 92 | -0.04 | 0.57 | -0.07 | -1.36 | 2.02 |
| Maternal Prosody Composite Reunion Min 2 | 88 | -0.10 | 0.60 | -0.18 | -1.89 | 2.00 |
| *Infant Behavioral Distress (Raw values)* |  |  |  |  |  |  |
| Still Face | 93 | 0.26 | 0.14 | 0.25 | 0.03 | 0.81 |
| Reunion Min 1 | 93 | 0.16 | 0.12 | 0.14 | 0.00 | 0.71 |
| Reunion Min 2 | 93 | 0.20 | 0.17 | 0.15 | 0.00 | 0.77 |
| SF to Reunion Change | 93 | -0.06 | 0.20 | -0.04 | -0.57 | 0.60 |
| *Infant Behavioral Distress (Normalized z scores)* |  |  |  |  |  |  |
| Still Face | 93 | 0.42 | 0.81 | 0.45 | -1.51 | 2.92 |
| Reunion Min 1 | 93 | -0.29 | 0.89 | -0.32 | -2.55 | 2.24 |
| Reunion Min 2 | 93 | -0.11 | 1.14 | -0.22 | -2.55 | 2.55 |
| Still Face to Reunion Change Score | 93 | -0.53 | 1.26 | -0.33 | -3.68 | 2.61 |
| *Infant Heart Rate* |  |  |  |  |  |  |
| Still Face | 88 | 149.80 | 11.61 | 147.83 | 126.47 | 179.26 |
| Reunion Min 1 | 87 | 146.18 | 13.90 | 146.30 | 115.88 | 191.19 |
| Reunion Min 2 | 83 | 146.14 | 13.38 | 144.66 | 116.43 | 178.68 |
| Still Face to Reunion Change Score | 83 | -2.98 | 7.83 | -2.28 | -23.53 | 20.16 |
| *Infant Respiratory Sinus Arrhythmia (RSA)* |  |  |  |  |  |  |
| Still Face | 88 | 3.17 | 1.00 | 3.08 | 1.13 | 5.74 |
| Reunion Min 1 | 87 | 3.34 | 1.18 | 3.27 | 0.62 | 7.09 |
| Reunion Min 2 | 83 | 3.33 | 1.04 | 3.08 | 1.36 | 5.73 |
| Still Face to Reunion Change Score | 83 | 0.18 | 0.65 | 0.06 | -1.46 | 2.84 |

**Table 2.** Paired t-test results for infants change from still face to second half of reunion.

| **Variable** | **Mean** | **95% CI** |  | **t** | **DF** | **p** | **Cohen's d** |
| --- | --- | --- | --- | --- | --- | --- | --- |
|  |  | **Low** | **High** |  |  |  |  |
| Behavioral distress | -.53 | -.79 | -.27 | -4.05 | 92 | <.001 | .42 |
| Heart rate | -2.98 | -4.69 | -1.27 | -3.46 | 82 | .001 | .38 |
| Respiratory sinus arrhythmia | .18 | .03 | .32 | 2.47 | 82 | .016 | .27 |

Bivariate correlations are presented in Table 3. Infants’ behavioral distress during the still face episode and reunion was positively correlated. In addition, infants decreased their behavioral distress more in reunion if they had been more distressed during still face (*r* = -.45).

**Table 3.** Correlation matrix for key study variables measured during Still Face (SF) and Reunion. Reunion infant measures were binned into 1-minute intervals to examine stability and change over the course of resumption of co-regulation between infant and mother. Change scores were calculated as the difference between Still Face and end of reunion (minute 2).

| Variable | 1 | 2 | 3 | 4 | 5 | 6 | 7 | 8 | 9 | 10 | 11 | 12 | 13 |
| --- | --- | --- | --- | --- | --- | --- | --- | --- | --- | --- | --- | --- | --- |
| *Maternal Prosody Composite* |  |  |  |  |  |  |  |  |  |  |  |  |  |
| 1. Reunion Min 1 |  |  |  |  |  |  |  |  |  |  |  |  |  |
| 2. Reunion Min 2 | .75** |  |  |  |  |  |  |  |  |  |  |  |  |
| *Infant Behavioral Distress* |  |  |  |  |  |  |  |  |  |  |  |  |  |
| 3. Still Face | .06 | .20 |  |  |  |  |  |  |  |  |  |  |  |
| 4. Reunion Min 1 | -.21* | -.17 | .35** |  |  |  |  |  |  |  |  |  |  |
| 5. Reunion Min 2 | -.21* | -.15 | .21* | .62** |  |  |  |  |  |  |  |  |  |
| 6. $\Delta$ SF to End of Reunion | -.23* | -.26* | -.45** | .34** | .78** | | | | | | | | |
| *Infant Heart Rate* |  |  |  |  |  |  |  |  |  |  |  |  |  |
| 7. Still Face | .04 | .25* | .11 | .11 | .16 | .07 |  |  |  |  |  |  |  |
| 8. Reunion Min 1 | -.01 | .13* | -.00 | .22* | .20 | .19 | .87** |  |  |  |  |  |  |
| 9. Reunion Min 2 | -.08 | .05 | -.12 | .11 | .30** | .35** | .81** | .88** |  |  |  |  |  |
| 10. $\Delta$ SF to End of Reunion | -.27* | -.24* | -.26* | .16 | .38** | .51** | -.05 | .24* | .54** | | | | |
| *Infant Respiratory Sinus Arrhythmia (RSA)* |  |  |  |  |  |  |  |  |  |  |  |  |  |
| 11. Still Face | .13 | -.03 | -.05 | .01 | -.00 | .03 | -.49** | -.40** | -.41** | .01 |  |  |  |
| 12. Reunion Min 1 | .11 | .03 | -.04 | -.01 | -.01 | .01 | -.29** | -.34** | -.30** | -.09 | .80** |  |  |
| 13. Reunion Min 2 | .16 | .11 | -.09 | -.02 | -.09 | -.03 | -.28** | -.28** | -.31** | -.13 | .79** | .86** |  |
| 14. $\Delta$ SF to End of Reunion | .09 | .19 | -.04 | -.01 | -.10 | -.07 | .26* | .15 | .10 | -.21 | -.20 | .22* | .44** |

\* $p < .05$ . \*\* $p < .01$ .

Infants’ heart rates were highly positively correlated among all observations (range: .81, .88) but there was no significant association between still face heart rate and the magnitude of change to the reunion. Infants’ RSA followed a similar pattern of high correlations among measurements (range: .79, .86) but did not have a significant correlation between still face level and change to reunion.

Infants’ heart rate and RSA were negatively correlated during all simultaneous measurements (range: -.31, -.49), but heart rate and RSA changes from still face to the end of reunion were not significantly correlated (*r* = -21). Additionally, there was a small positive correlation *(r* = .26) between infant heart rate during still face and RSA change scores, indicating that infants with higher heart rates during still face had greater increases in RSA as they moved into the reunion episode.

During reunion, behavioral distress was higher in infants with higher heart rates (minute 1, *r* = .22, minute 2, *r* = .30). However, heart rate was not correlated with behavioral distress during still face. Higher behavioral distress during still face was associated with greater reductions in heart rate from still face to the end (second minute) of the reunion episode (*r* = -.26). Changes in heart rate from still face to reunion were highly positively correlated with concurrent changes in behavioral distress, demonstrating that infants whose heart rate decreased the most exhibited the greatest decreases in behavioral distress (r = -.51). Unlike heart rate, infants’ RSA during still face and reunion episodes was not significantly related to any behavioral distress measures (bottom left of Table 3).

Finally, the maternal prosody composite was highly correlated during the first and second minutes of reunion (r = .75). Greater values on the maternal prosody composite during reunion was correlated with larger decreases in infants’ behavioral distress and heart rate (see first columns of Table 3). The maternal prosody composite during the first minute of reunion was also negatively correlated with lower overall levels of infants’ behavioral distress in both segments of the reunion.

### Infants’ biobehavioral state and maternal prosody

A series of regression models examined the relation between caregivers’ prosody composite during the beginning of reunion (minute 1) and changes in infants’ biobehavioral state during the Still Face episode. In each model, the outcome variable was change in infants’ behavioral distress or physiology from Still Face to the end of reunion (minute 2). Interactions between the maternal prosody composite and infants’ biobehavioral state were tested and the interaction terms were retained if the parameter achieved statistical significance at alpha = .05.

Table 4 displays results of the models. As illustrated in Figure 3, greater values on the maternal prosody composite were associated with a decrease in infants’ behavioral distress and heart rate. The interaction terms for these models were excluded using the criteria described above. However, there was a significant interaction between the maternal prosody composite and infants’ RSA during the Still Face episode. Infants who had low RSA during the Still Face stressor exhibited increased RSA during reunion when values on the maternal prosody composite were higher. However, infants whose RSA was already high demonstrated no association between RSA changes and the maternal prosody composite (Figure 3 panel c). To test whether the interaction could be explained by individual differences in infant baseline RSA, an additional model was tested in which baseline RSA collected prior to mother-infant play (see Methods) was included as an interaction term. Since this added variable was not significant and did not change the maternal prosody composite by infant SF RSA interaction, the analysis is consistent with the effect being specific to change from the challenge to the reunion and not dependent on individual differences in baseline RSA. Comparison follow-up analyses using individual indicators showed that effects were strongest and most consistent for the combined prosody composite, though there was some evidence that modulation depth was particularly associated with decreases in infant heart rate and mid-frequency power was particularly related to infant RSA (Supplemental Material Figure S3).

**Table 4.** Model results predicting change in infant behavioral distress, heart rate, and respiratory sinus arrhythmia (RSA) from Still-Face to the second half of Reunion. Interaction terms for behavioral distress and heart rate were non-significant at alpha = .05 and thus excluded from final models.

| **Parameter** | **B** | **SE** | **t** | **p** |
| --- | --- | --- | --- | --- |
| *Outcome: Change in Infant Behavioral Distress* |  |  |  |  |
| Intercept | -0.513 | 0.125 | -4.104 | 0.000 |
| Maternal Prosody Composite Min 1 | -0.492 | 0.218 | -2.253 | 0.027 |
| *Outcome: Change in Infant Heart Rate* |  |  |  |  |
| Intercept | -2.918 | 0.826 | -3.532 | 0.001 |
| Maternal Prosody Composite Min 1 | -3.532 | 1.423 | -2.482 | 0.015 |
| *Outcome: Change in Infant Respiratory Sinus Arrhythmia (RSA)* |  |  |  |  |
| Intercept | 0.722 | 0.237 | 3.051 | 0.003 |
| Maternal Prosody Composite | 1.050 | 0.468 | 2.244 | 0.028 |
| Infant RSA (Still Face episode) | -0.171 | 0.072 | -2.393 | 0.019 |
| Maternal Prosody Composite X Infant SF RSA (Interaction Term) | -0.280 | 0.138 | -2.033 | 0.045 |

**Figure 3.**
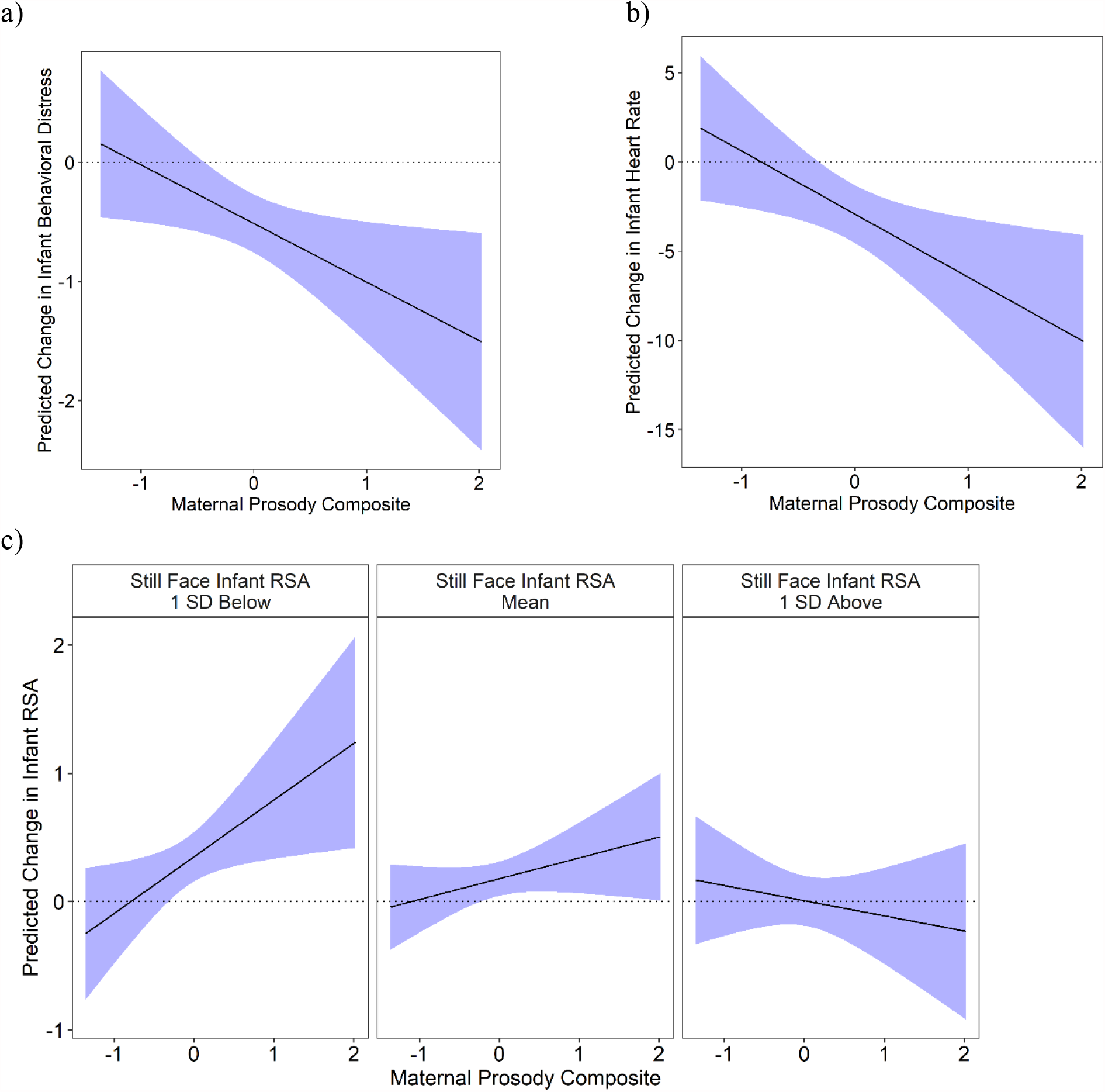
Predicted values and 95% confidence intervals for changes in infant biobehavioral state. Model predicted values are visualized for behavioral distress (a) and infant heart rate (b) from Still Face to the second half Reunion as a function of the maternal prosody composite. High values on the maternal prosody composite were associated with decreases in both infant behavioral distress and heart rate. Interactions between the maternal prosody composite and infant state at SF were not significant in either model. The bottom panel (c) depicts changes in infant respiratory sinus arrhythmia from the Still Face episode to the second half of reunion as a function of the maternal prosody composite. High maternal prosody composite scores were related to increases in cardiac vagal tone in infants who had low respiratory sinus arrhythmia (RSA) during still face.

Cross-lagged panel models were used to examine temporal dynamics of the maternal prosody composite and infant state. The cross-lagged panel model for infant heart rate showed that both infant heart rate and the maternal prosody composite were highly stable throughout reunion and that infant heart rate during Still Face did not directly predict the maternal prosody composite at the beginning of reunion (Figure 4). However, there was evidence that maternal prosody composite and infant heart rate both predicted changes in the other variable over the course of reunion. Higher infant heart rate during the first minute of reunion predicted increased values on the maternal prosody composite during the second minute while greater maternal prosody composite values during the first minute predicted a decrease in infant heart rate during the second minute.

**Figure 4.**
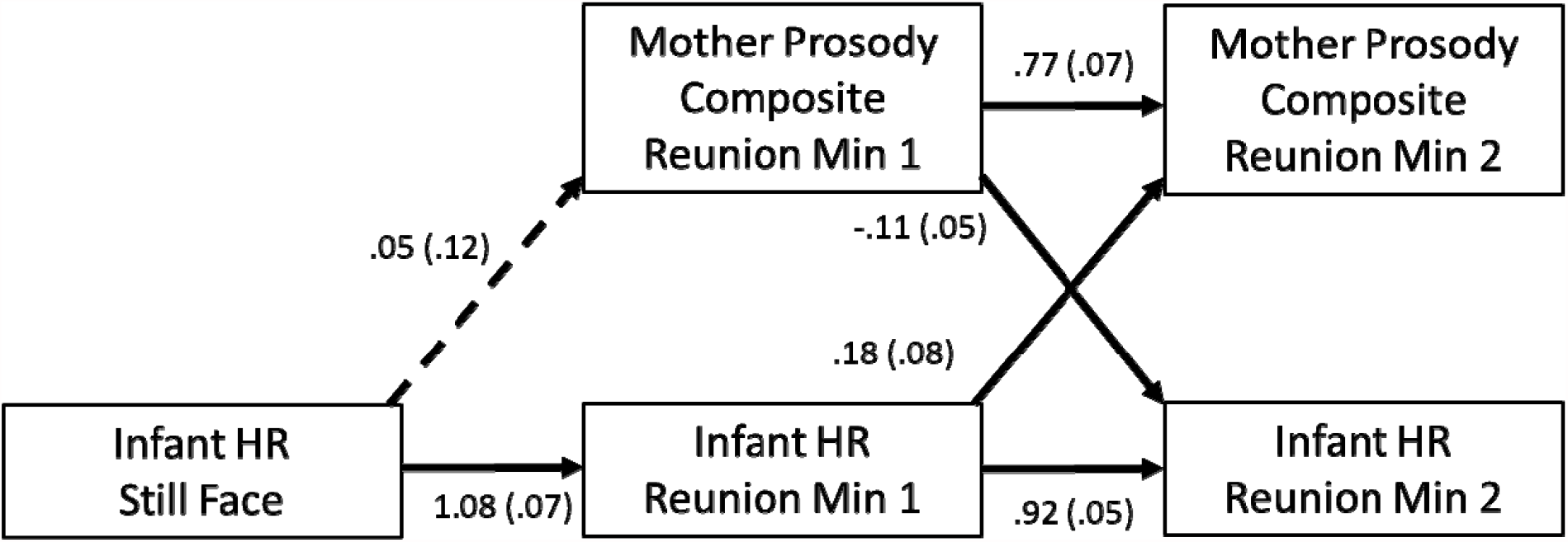
Cross-lagged panel model of infant heart rate (HR) and the maternal prosody composite over the course of Still Face and reunion episode in 1-minute intervals. Tested paths are presented with standardized point estimates and standard errors (in parentheses). Solid line arrows represent paths whose p values < .05 and dotted lines those whose p values >= .05. Model fit indices: Chi-Square = 8.266, degrees of freedom = 4, Tucker-Lewis Index = .97, Comparative Fit Index = .99, Root-Mean-Square Error of Approximation = .107 [90% CI: .000,.210]

The cross-lagged panel model for infant behavioral distress and the maternal prosody copmosite resulted in a poor fit. Modification indices revealed that fit could be improved by including a covariance between the maternal prosody composite and infant behavioral distress at minute 1 (Figure 5). This model showed that the infant behavioral distress variable was less stable than infant physiological variables. While mother and infant variables at minute 1 did not predict the other at minute 2, there was a significant negative covariance at minute 1 between the variables.

**Figure 5.**
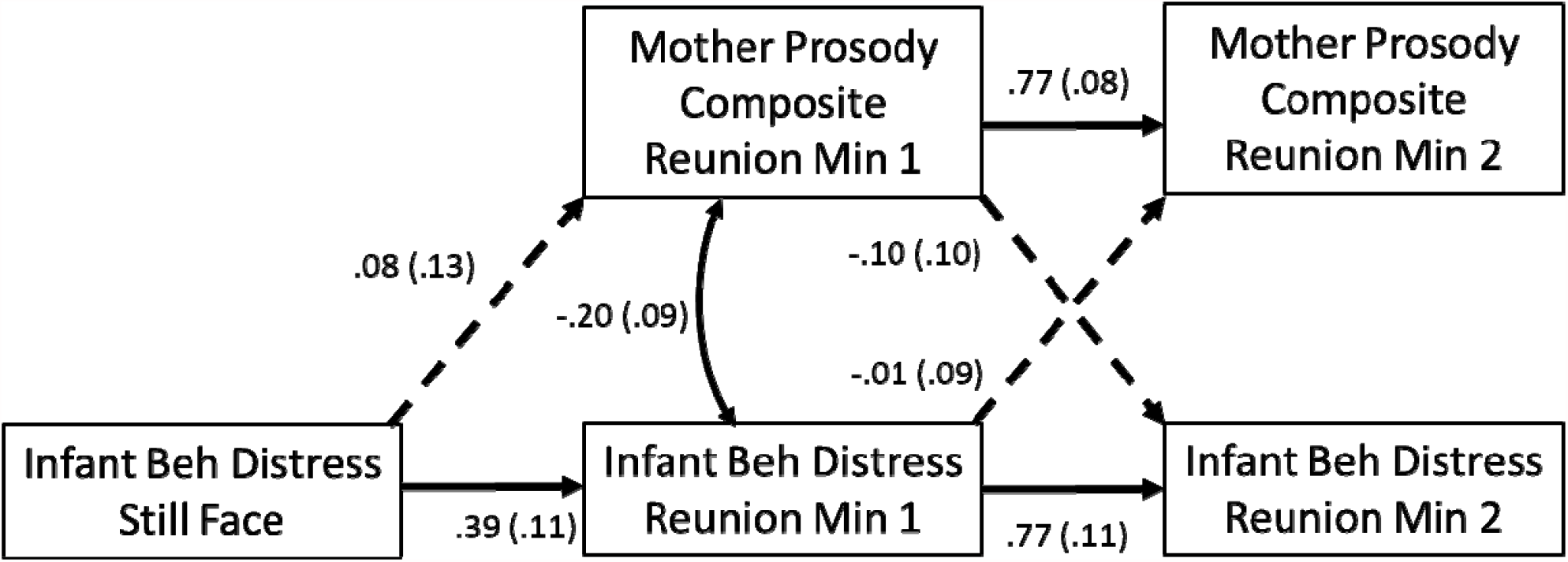
Cross-lagged panel model of infant behavior distress and the maternal prosody composite over the course of Still Face and reunion episode in 1-minute intervals. Tested paths are presented with standardized point estimates and standard errors (in parentheses). Solid line arrows represent paths whose p values < .05 and dotted lines those whose p values >= .05. Model fit indices: Chi-Square = 4.37, degrees of freedom = 2, Tucker-Lewis Index = .91, Comparative Fit Index = .98, Root-Mean-Square Error of Approximation = .113 [90% CI: .000, .260]

Due to the observed interaction between infant Still Face RSA and the maternal prosody composite, modeling of the temporal dynamics of this variable was not possible as planned. Attempts to fit the model without an interaction supported this, showing unacceptable fit that could not be improved by simple adjustments suggested by modification indices (Chi-Square = 16.275, degrees of freedom = 4, TLI = .89, CFI = .96, RMSEA = .196 [90% CI: .103, .299]). There was an insufficient number of observations and degrees of freedom to adequately model this complex interaction. Because poorly fitting models do not provide credible parameter estimates, the model results are not presented or discussed.

## Discussion

This study demonstrates that maternal prosodic features - defined by strong power in the frequency band of perceptual advantage (500-5000Hz), weak high frequency power (5000 Hz +), and spectro-temporal modulation – are associated with biobehavioral calming in infants following a social stressor. This is consistent with a model proposing that a combination of frequency band and frequency modulation prosodic features may provide cues of safety that would facilitate the infant’s state regulation (Kolacz et al., 2018; Porges & Lewis, 2010). In addition, the results demonstrate that infant physiological regulation as measured by heart rate and maternal prosody are mutually co-regulated across time, with infant heart rates decreasing after interaction with mothers with high prosody and mothers’ prosody increasing when their infants’ heart rates were high. While the associations of prosody with heart rate and behavioral distress were consistent at the group level, only infants with lower cardiac vagal tone increased their cardiac vagal tone when paired with mothers with highly prosodic features. This suggests that infants whose cardiac vagal tone is low (indicating low parasympathetic activity) may benefit most from the prosodic voices of their caregivers.

Maternal prosody, when assessed over 1-minute intervals, showed stability during reunion (standardized coefficient = .77 in the cross-lagged panel models), although it is unknown whether this level reflects a stable characteristic of dyads or the individual mothers beyond the observed interaction. However, mothers also varied in their prosodic features over time, and these features were related to changes in infant biobehavioral state, supporting the possibility that caregivers and infants were co-regulating by responding to one another’s states. These results highlight the importance of considering dynamic changes that may be masked when averaging behavior across entire observation periods (e.g. such as entire phases during the Still Face; Ekas, Haltigan, & Messinger, 2013).

Heart rate serves as a functional metric of broad infant autonomic state that includes both parasympathetic and sympathetic influences. Our findings demonstrate that infants’ autonomic state may be modulated by maternal prosody, with mothers’ high prosodic features (strong mid-frequency power and spectro-temporal modulation) likely providing cues for infant autonomic soothing. This underscores how infant regulation unfolds across the reunion phase, corroborating recently reported timescales of arousal and distress modulation (Abney, daSilva, & Bertenthal, 2021; Cirelli & Trehub, 2020) and aligning with observations that mothers adjust their acoustic features depending on observed infant states (Smith & Trainor, 2010; Kitamura & Burnham, 2003; Stern, Spieker, Barnett, & MacKain, 1983).

Furthermore, infants’ heart rates during the first minute of reunion predicted increases in maternal prosody in the second minute of reunion. Though caregivers do not have direct knowledge of their infants’ heart rates, changes in autonomic state are integrated with social expression as part of the social engagement system (Porges, 1995, 2001, 2007). Although there are intrinsic individual differences in HR, such as differences in the function of the heart’s internal pacemaker and maturing of the ANS, that make associations of *between*-individual correlations of behavior and heart rate weak, *within*-individual associations between HR and behavioral changes can illuminate the strength of the link between behavior and physiology. In our study, there were moderate correlations between heart rate and behavioral distress changes from Still Face to the end of reunion (r = .51), which are comparable to those reported in other studies (Conradt & Ablow, 2010; Ham & Tronick, 2006). Such outward behavioral reflections of internal state, including facial expression and acoustic features, may elicit changes in caregiver vocal prosody (e.g. Galvez-Pol, Antoine, Li, & Kilner, 2020). The increased maternal prosody in dyads in which infants had high heart rates may function as a compensatory maternal response to help the infant regulate autonomic state when the infant enters distress.

Mothers’ own biobehavioral state and subjective distress ratings are impacted by infants’ acoustic features, with higher infant cry frequencies and greater pitch instability eliciting greater reports of maternal subjective distress, urgency, aversion, and discomfort as well as stronger sympathetic responses (Gustafson & Green, 1989; Zeskind & Marshall, 1988). Although the interpretation of our data is focused on aggregated patterns of maternal prosody in relation to infant states, it is possible that shifts in caregiver prosody reflect an individual difference and could provide a measure of caregiver sensitivity. Caregiver voices may also reflect their unique contributions to children’s state regulation. For instance, Palestinian mothers with depressive symptoms and history of war trauma are rated by coders as having more negative and lower positive emotion when singing to their infants and their expression of fear and tension are associated with lower infant positive affectivity (Punamäki, Vänskä, Quota, Perko, & Diab, 2020).

Infant cardiac vagal tone did not have strong relations with heart rate or behavioral distress, suggesting that in this sample biobehavioral state was not strongly regulated by a withdrawal of the vagal brake (Porges, Doussard-Roosevelt, Portales, & Greenspan, 1996). This may be a feature of the young sample in this study, as the cardiac vagal pathways mature over the first months of life (Porges & Furman, 2011) and respiratory sinus arrhythmia increases rapidly over this time (Alkon, Boyce, Davis, & Eskenazi, 2011; Bornstein & Suess, 2000). While temporal dynamics could not be modeled with the study data due to interaction, the results suggest maternal prosody may be especially potent for regulating state when infant cardiac vagal tone is low. This is consistent with the neurophysiological linkages between the ventral vagal complex, which gives rise to vagal tone, and the social engagement system that is posited by polyvagal theory. When physiological state regulation via the social engagement system is low, greater maternal prosody may provide social cues of safety that activate the cardiac vagal brake to calm infant state. In our data, this effect was not explained by baseline levels of cardiac vagal tone, suggesting that this reflects a response to the still face paradigm, but additional studies are needed to examine this further.

These results also align with recent research findings that infants with low vagal tone rely more on their mothers as an external source of support to co-regulate their behavior compared to infants with higher vagal tone, who may have stronger capacity for self-regulation. For example, Busuito, Quigley, Moore, Voegtline, & DiPietro (2019) found that lower levels of infant vagal tone were associated with higher behavioral synchrony with their mothers; the authors interpreted these findings to reflect that low vagal tone infants may have required greater synchoronous engagement to facilitate their biobehavioral regulation across the FFSF. Toddlers with low cardiac vagal tone benefit more from caregiver interaction with regard to exhibiting social approach (Grady & Callan, 2019) and executive function ability (Gueron-Sela et al., 2017) compared to their peers with high cardiac vagal tone. Perry and colleagues found a relationship between maternal socialization behaviors and children’s emotion regulation only for children with lower levels of vagal suppression (Perry, Calkins, Nelson, Leerkes, & Marcovitch, 2012). Collectively, these studies underscore how maternal interaction may differentially contribute to the development of emotion regulation and executive function for children who have more difficulty with physiological regulation.

A related consideration is the extent to which mother’s own physiological regulation may co-vary with prosodic cues and interact with infants’ biobehavioral regulation. Maternal vagal regulation during infant negative affect has been associated with various forms of maternal sensitivity (e.g Leerkes, Su, Calkins, Supple, & O’Brien, 2016; Mills-Koonce et al., 2007; Moore et al., 2009), raising the possibility that a similar mechanism may operate vis-a-vis maternal prosody. Given that the still-face is a stressor for the mother as well as the infant (Mayes, Carter, Egger, & Pajer, 1991), it is reasonable to expect that the mother’s own regulatory capacity will contribute to her ability to co-regulate her infant’s biobehavioral distress.

Infants’ behavioral distress was less temporally stable than heart rate and cardiac vagal tone. When viewed in light of the significant covariance of mother prosody and infant behavioral distress at reunion minute 1, it is possible that these temporal dynamics are unfolding over a faster time scale than could be measured with the current study data. Although directionality cannot be established, this covariance was negative, suggesting that greater maternal prosody may provide cues for reduction of infants’ distress or that infant behavioral distress may induce decreases in maternal prosodic features. This analysis focused on changes within 1-minute bins to provide enough data for aggregation. Measurement of maternal voice during naturalistic observation generates sparse data – data is not available when mothers are silent or their vocalizations overlap with infant noises. Though not optimal for micro-scale analyses, the data presented here may inform future studies where protocol design and equipment permits richer data streams that can capture prosody dynamics at faster time scales. Future studies are needed to examine the time scales at which caregiver prosody and infant state regulation coordinate, using methods that optimize fine-grained vocal data collection without diminishing the ecological validity of the mother-infant interaction.

Though the maternal prosody composite was consistently associated with biobehavioral regulation infants, there was some evidence that individual acoustic components may be particularly salient for physiological infant state changes, with modulation depth having the strongest association with decreases in infants’ heart rate and mid-frequency power being most associated with RSA increases in infants with low RSA. However, given the largely consistent direction of effects of the prosody components in predicting infant changes, additional studies are needed to determine whether this effect can be replicated.

## Limitations

Cross-lagged panel models were used in this study to examine temporal dynamics of maternal prosody and infants’ biobehavioral regulation. While these models can reveal time-related dynamics in aggregated data, the results include a combination of within- and between-dyad effects (Berry & Willoughby, 2017). Future studies with larger samples may be able to disaggregate these effects and distinguish between these two sources of variance.

Future studies should also examine the effects of maternal prosody with a more diverse sample or with samples that include infants at-risk (e.g. Boeve, Beeghly, Stacks, Manning, & Thomason, 2019; Conradt & Ablow, 2010). A recent meta-analysis found that at-risk infants generally do not show RSA increases from still-face to reunion (Jones-Mason et al., 2018). Extending our finding that infants with low vagal tone showed greater RSA increases when mothers had more prosodic acoustic features, it is reasonable to expect that maternal prosody would support physiological regulation for at-risk infants who may have regulatory difficulties. Moreover, other research suggests that child-directed speech varies as a function of socioeconomic status and education level (e.g Rowe, 2008), raising the possibility that maternal prosody may function similarly.

In addition, this study focused specifically on caregiver prosody. Maternal cues that regulate infants’ state are likely multi-modal and may include facial expressions, touch, and gesture (Chong, Werker, Russell, & Carroll, 2003; Stack & Arnold, 1998). In this study, mothers were explicitly prohibited from touching their infants (Feldman et al., 2010), but the mothers’ facial expression and gestures could not be ruled out. Additional research is needed to isolate the effects of the mothers’ voice from other modalities of social communication to examine whether the effects of the mother on the infants’ state are directly mediated by prosodic features.

Although this analysis was limited to maternal speech given its predominance during the Still Face paradigm, more data are needed to assess how the acoustic features of the full repertoire of maternal vocalizations relate to infant biobehavioral state. For instance, post hoc analyses showed that mothers who sang during the reunion had infants with higher RSA and lower HR during the course of the entire protocol, including baseline prior to play. This broad distinction in the present study prevents conclusions about whether the mothers’ singing is the distinguishing factor or whether some other type of features (e.g., emotional attunement in the dyad) is associated with a propensity to sing. However, since singing is known to be an efficient method of regulating physiological infant state (Cirelli & Trehub, 2020), this suggests more work is needed to understand mechanisms of infant regulation that distinguish dyads in which mothers spontaneously sing.

## Conclusion

The present study is the first to use a novel, theoretically-informed composite measure of prosody based on the frequency bands and modulation of voice to study the naturalistic dynamics of caregiver vocal features and infant biobehavioral state. The results, when taken together, suggest that infant and caregiver co-regulate state after a disruptive challenge, with maternal prosody levels adjusting depending on infant state and, in turn, predicting decreases in infant biobehavioral distress. These findings expand the conceptualizations of the mechanism of dyadic emotional regulation in infants’ early life, pointing to new research and potential intervention opportunities for caregivers and infants.

