## Supplemental Materials for "Associations between Acoustic Features of Maternal Speech and Infants’ Emotion Regulation following a Social Stressor"

Jacek Kolacz^1^ *

Elizabeth B. daSilva^2^ *

Gregory F. Lewis^1, 3^

Bennett I. Bertenthal^4^

Stephen W. Porges^1,5^

^1^Traumatic Stress Research Consortium, Kinsey Institute, Indiana University

^2^Indiana University-Purdue University Columbus, Columbus, IN

^3^Intelligent Systems Engineering, Indiana University

^4^Department of Psychological and Brain Sciences, Indiana University, Bloomington, IN

^5^Department of Psychiatry, University of North Carolina at Chapel Hill

*JK and EBD should be considered joint first author

Table S1. Correlations between speech-only and all-vocalization audio files. In order to conduct sensitivity analysis, files were manually re-edited to capture all maternal vocalizations. While the original analysis included only voiced speech (see main text), the re-analysis also permitted singing, whispering, rhythmic chanting, and any other mouth-originating sounds that the mother made. The expanded criteria added a median of 6.32 seconds to audio included for the analysis from the 2-minute reunion (median added to first minute = 1.58 seconds, median added to second minute = 5.13 seconds; medians do not simply sum to the total for 2 minutes due to some missing data in each portion of the reunion). Correlations between acoustic variables derived from the two editing strategies were highly correlated.

|  | Reunion  Minute 1 | Reunion  Minute 2 |
| --- | --- | --- |
| Modulation depth | .76 | .76 |
| High frequency power | .80 | .72 |
| Mid frequency power | .88 | .82 |
| Prosody composite | .89 | .84 |

Table S2. Sensitivity analysis using all maternal vocalizations to calculate the prosody composite. The re-calculated prosody values were used to predict change in infant behavioral distress, heart rate, and respiratory sinus arrhythmia (RSA) from Still-Face to the second half of Reunion. Consistent with table 4 (maternal speech only models), interaction terms for behavioral distress and heart rate were non-significant at alpha = .05. However, compared to table 4 (speech only), there are no significant effects. Direct effect model predicting RSA from the prosody composite only was also non-significant (prosody *B* = .099, *SE* = .141, *p* = .484).

| **Parameter** | **B** | **SE** | **t** | **p** |
| --- | --- | --- | --- | --- |
| *Outcome: Change in Infant Behavioral Distress* |  |  |  |  |
| Intercept | -0.536 | 0.130 | -4.122 | 0.000 |
| Maternal Prosody Composite (all vocalizations) | -0.316 | 0.262 | -1.206 | 0.231 |
| *Outcome: Change in Infant Heart Rate* |  |  |  |  |
| Intercept | -3.060 | 0.861 | -3.553 | 0.001 |
| Maternal Prosody Composite (all vocalizations) | -1.913 | 1.686 | -1.134 | 0.260 |
| *Outcome: Change in Infant Respiratory Sinus Arrhythmia (RSA)* |  |  |  |  |
| Intercept | 0.695 | 0.250 | 2.784 | 0.007 |
| Maternal Prosody Composite (all vocalizations) | 0.952 | 0.661 | 1.441 | 0.154 |
| Infant Cardiac Vagal Tone Still Face | -0.160 | 0.075 | -2.119 | 0.037 |
| Maternal Prosody Composite X Infant SF RSA | -0.256 | 0.198 | -1.291 | 0.200 |

Figure S1. Cross-lagged panel models of associations between the maternal prosody composite (all vocalizations) with: a) infant behavior distress and b) heart rate with over the course of Still Face and reunion episode in 1-minute intervals. Tested paths are presented with standardized point estimates and standard errors (in parentheses). Solid line arrows represent paths whose p values < .05 and dotted lines those whose p values >= .05. In the behavioral distress model, the covariance that was required for model fit with the speech-only vocalization data (Figure 5 in main text) was not significant within this sensitivity analysis. Because that covariance made no substantial contribution to model fit that parameter was excluded.

Behavioral distress model fit indices: Chi-Square = 3.00, degrees of freedom = 4, Tucker-Lewis Index = 1.00, Comparative Fit Index = 1.00, Root-Mean-Square Error of Approximation = .000 [90% CI: .000, .137]

Heart rate model fit indices: Chi-Square = 6.003, degrees of freedom = 4, Tucker-Lewis Index = .98, Comparative Fit Index = .99, Root-Mean-Square Error of Approximation = .073 [90% CI: .000, .185]

a)

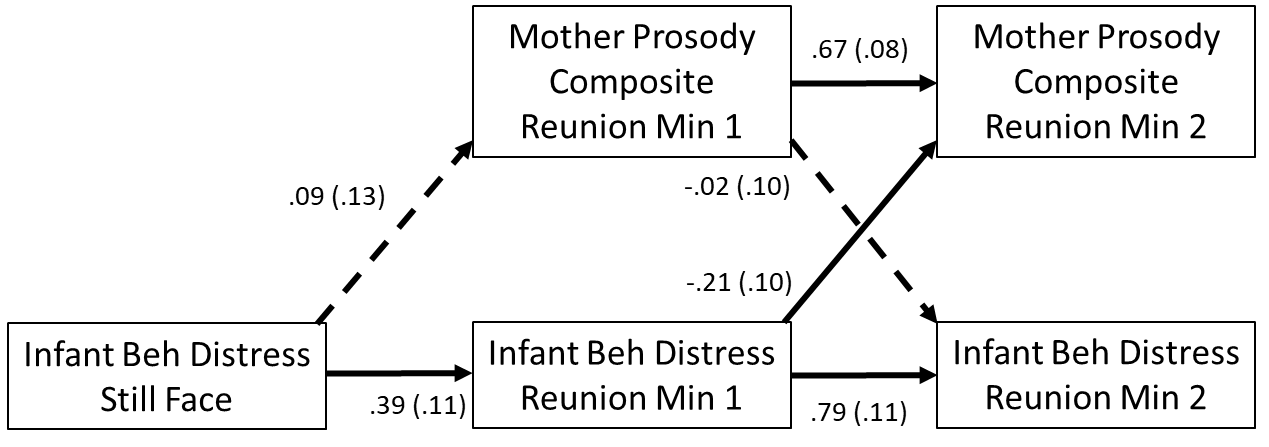

b)

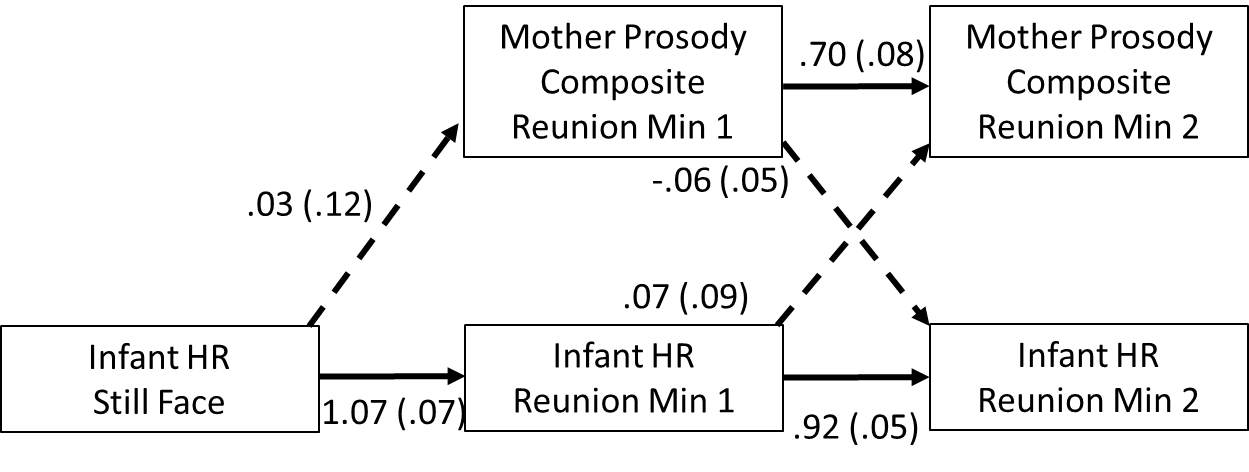

Figure S2. Fifteen out of 94 mothers sang during the 2-minute reunion after the Still Face (16%). The total duration time and low rate of spontaneous singing prevented analysis on whether singing vocalizations differed from non-singing vocalizations on the mother prosody composite and infant state variables. However, analyses were conducted to examine whether dyads in which mothers sang at any point during the post-Still Face reunion differed on key study variables. There was no evidence that mothers who sang differed on prosody metrics compared to non-signing mothers or that singing moderated effects of the prosody composite on infant HR, RSA, or behavioral distress from Still Face to reunion (all p > .25). However, infants whose mothers sang decreased more in distress between Still Face and the end of reunion (t(91) = 2.58, p = .011, No singing M = -.39, SD = 1.21; Singing M = -1.27, SD = 1.29; Cohen’s d = .71, see panel a below), though they did not significantly differ in absolute levels of distress in any phase of the Face-to-Face Still Face protocol. Patterns of infant HR and RSA showed strong evidence that mothers’ singing may be related to general organizational properties of individual dyads. Maternal signing was associated with higher RSA and lower HR across all aspects of the protocol, even during baseline when mother and infant were not interacting (panels b & c HR B = -9.83 95% CI: -17.16, -2.58 estimates from a repeated measures mixed effects model; RSA B = .69, 95% CI .03:1.34, estimates from a repeated measures model, no interactions between singing and protocol section were significant).

a)

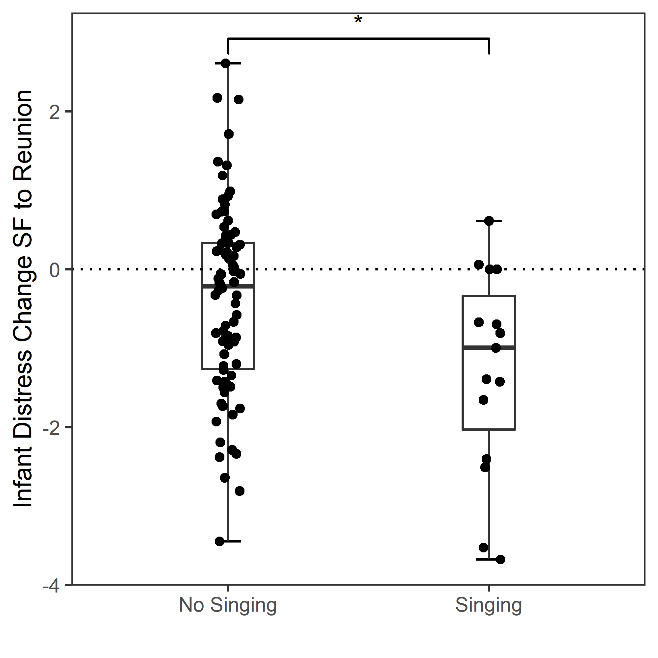

b)

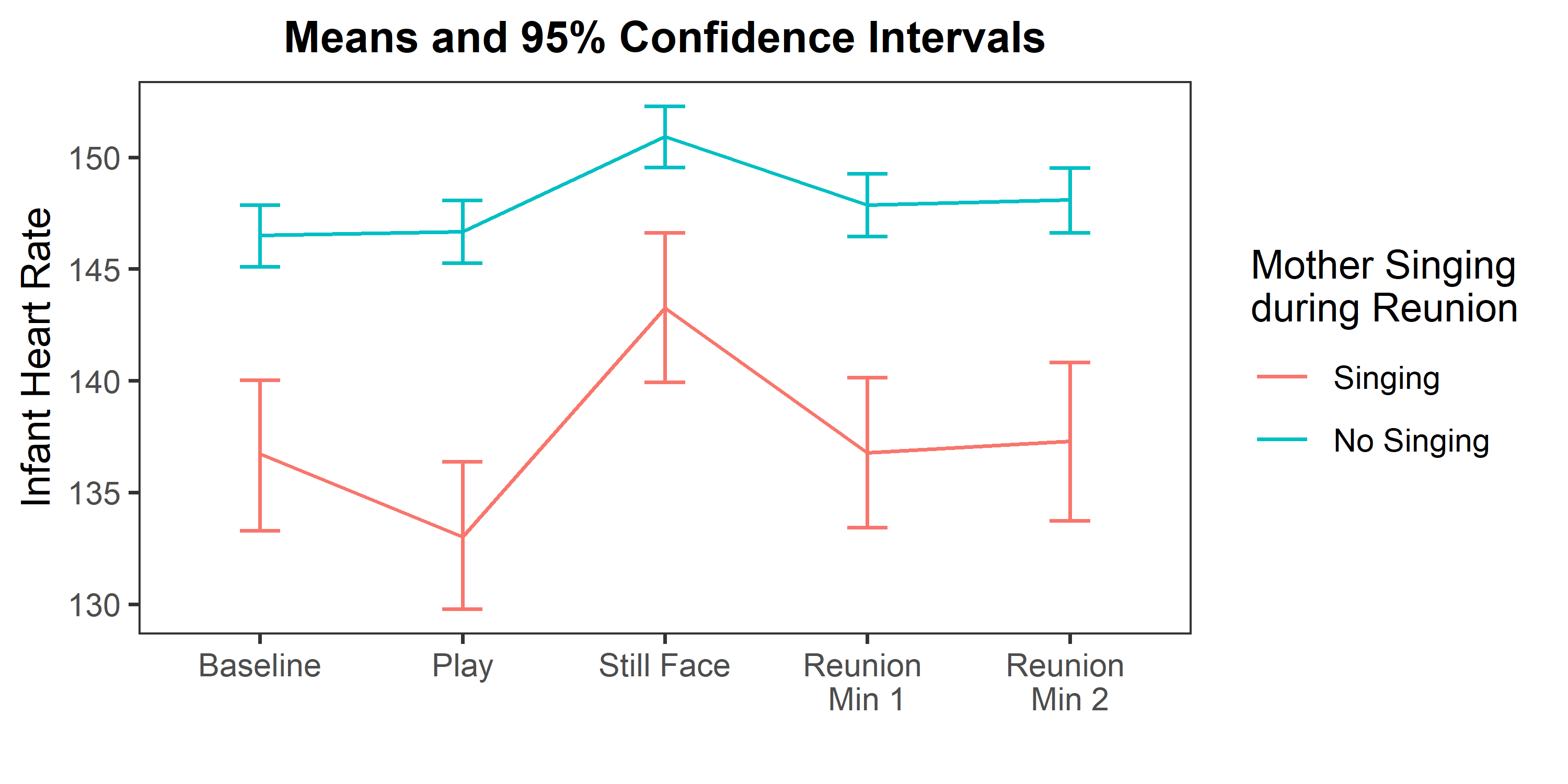

c)

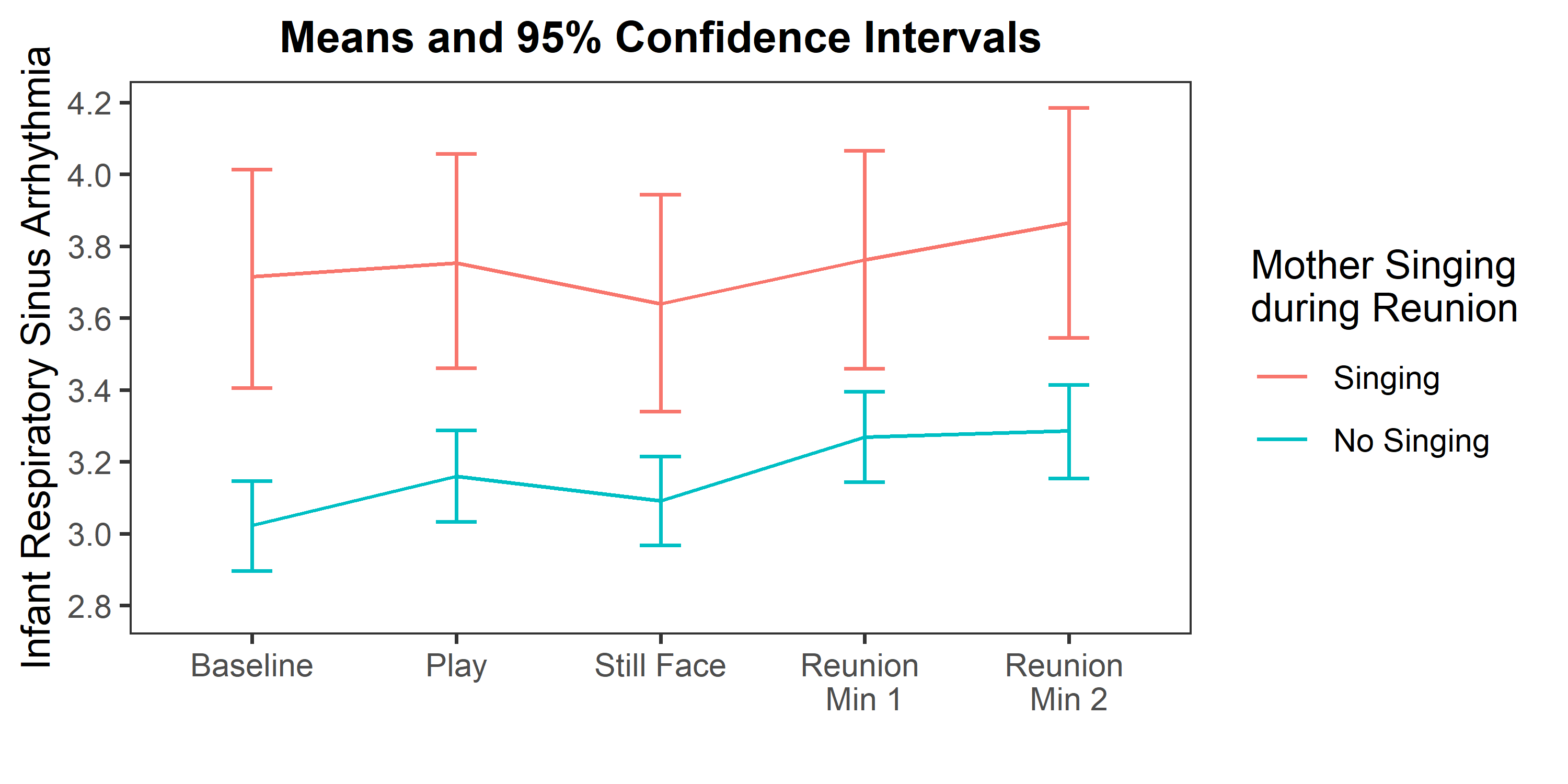

Figure S3. Individual components of the maternal prosody composite were each assessed for their association with changes in infant biobehavioral state measures from Still Face to the end of Reunion (2^nd^ minute). Regression models based on those reported in the main text (Figure 3 and Table 4) were repeated with individual prosody composite components (high frequency power, mid frequency power, and modulation depth). In the figure below, the inverse of high frequency power was used to correspond to composite scoring where lower high frequency power was treated as indicating more prosodic features. Few of the individual composite measures were significant predictors in isolation. Change in infant behavioral distress was not significantly predicted by any of the individual indicators, though effects were in a consistent direction with the composite measure. Infant decrease in heart rate was only significantly predicted by greater maternal modulation depth. Like the composite, none of the indicators had a direct effect on change in RSA. Prediction for change in RSA was additionally calculated at 1 SD below the mean during Still Face to correspond with effect reported with composite in the main analysis. Increase in RSA among infants with low RSA during Still Face was significantly predicted only by mid-frequency power. In sum, the prosody composite provided consistent predictions while individual indicators were rarely significant predictors in isolation, though all tended in the same direction as the composite. Sensitivity analyses including all maternal vocalizations (in contrast to speech displayed in the figure) showed a similar pattern of effects.

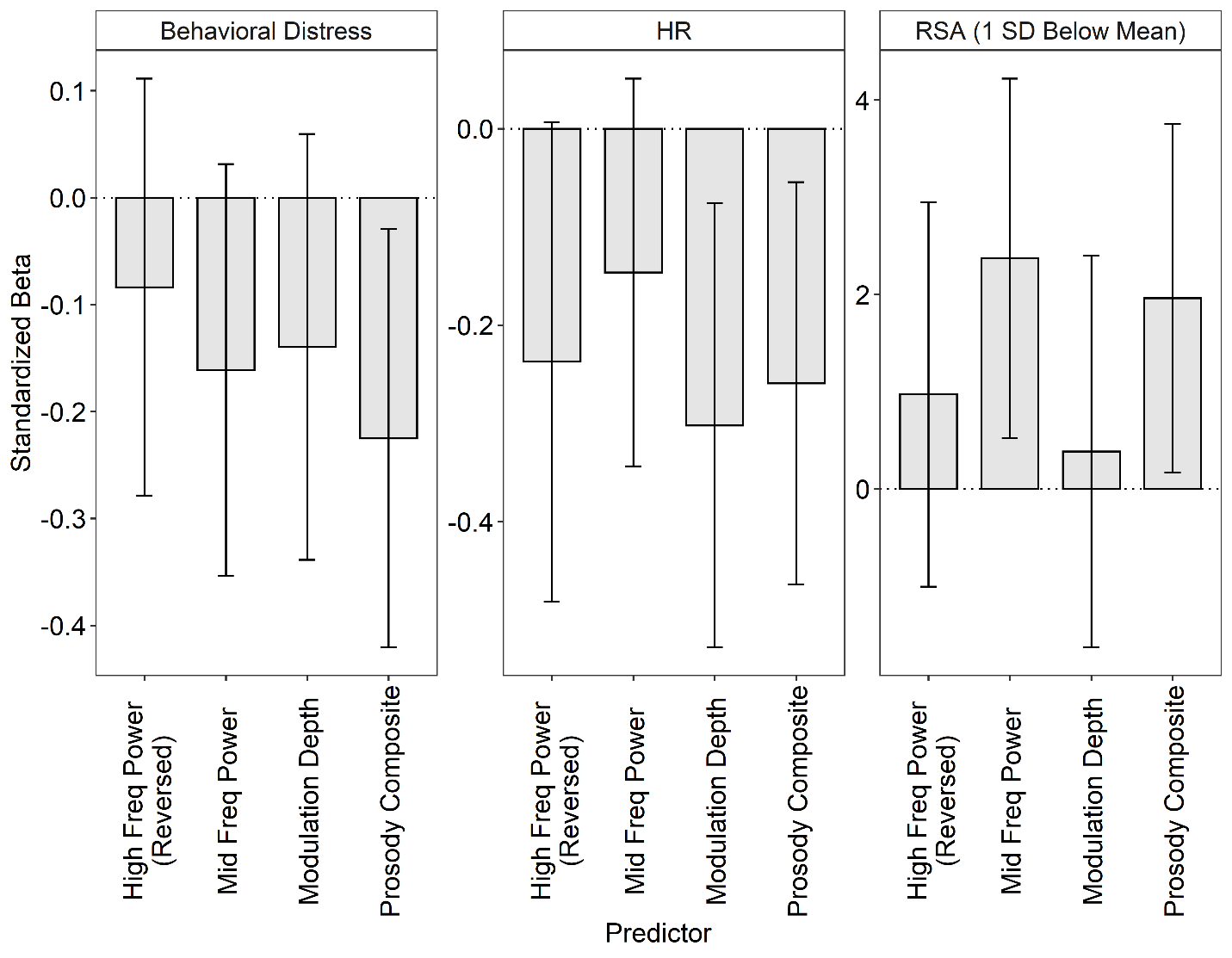
